# CRISPR-Cas13d as a molecular tool to achieve targeted gene expression knockdown in chick embryos

**DOI:** 10.1101/2024.08.03.606488

**Authors:** Minyoung Kim, Erica J. Hutchins

## Abstract

The chick embryo is a classical model system commonly used in developmental biology due to its amenability to gene perturbation experiments. Pairing this powerful model organism with cutting-edge technology can significantly expand the range of experiments that can be performed. Recently, the CRISPR-Cas13d system has been successfully adapted for use in zebrafish, medaka, killifish, and mouse embryos to achieve targeted gene expression knockdown. Despite its success in other animal models, no prior study has explored the potential of CRISPR-Cas13d in the chick. Here, we present an adaptation of the CRISPR-Cas13d system to achieve targeted gene expression knockdown in the chick embryo. As proof-of-principle, we demonstrate the knockdown of PAX7, an early neural crest marker. Application of this adapted CRISPR-Cas13d technique resulted in effective knockdown of PAX7 expression and function, comparable to knockdown achieved by translation-blocking morpholino. CRISPR-Cas13d complements preexisting knockdown tools such as CRISPR-Cas9 and morpholinos, thereby expanding the experimental potential and versatility of the chick model system.

## 1. INTRODUCTION

The chicken (*Gallus gallus*) is a classical model system in embryology research, providing foundational discoveries that underlie much of our understanding of developmental biology (Needham, 1959; Stern, 2005). The chick embryo, as an amniote, not only develops morphologically similarly to human embryos at comparable stages but also shares significant genomic sequence and gene function homology, making it an excellent organism for studying vertebrate development. Additionally, chick embryos develop externally, allowing for genetic manipulations at early stages of development (Mok et al., 2015). Pairing this model organism with cutting-edge technology can significantly extend the range of experiments that can be performed, as well as expand our knowledge of processes underlying embryogenesis.

A key strategy for understanding gene function during embryonic development involves the use of loss-of-function genetic tools that can reduce or ablate gene expression in model organisms. In the chick embryo, perturbing gene expression is possible with CRISPR-Cas9 (Gandhi et al., 2021; Gandhi et al., 2017) and morpholino oligonucleotides (Corey and Abrams, 2001). While morpholinos, which are available as splice- or translation-blocking antisense oligonucleotides, are highly effective in the avian embryo as a knockdown tool (Chacon and Rogers, 2019; Hutchins and Bronner, 2018, 2019; Hutchins et al., 2022; Kerosuo and Bronner, 2016; Manohar et al., 2020; Piacentino and Bronner, 2018), these reagents cannot be spatiotemporally restricted without targeted electroporations or direct injection (which is not always feasible) and can be cost-prohibitive for some research budgets as they must be synthesized commercially. CRISPR-Cas9 is also an effective, versatile knockdown tool in the chick that can be electroporated as a ribonucleoprotein complex (recombinant Cas9 protein in complex with *in vitro* transcribed guide RNA) (Gandhi et al., 2020; Hutchins and Bronner, 2018), or delivered as plasmid(s) (Gandhi et al., 2021; Gandhi et al., 2017) which enable spatially or temporally restricted knockouts with the use of different promoters. However, CRISPR-Cas9 plasmid-mediated knockout, because it functions at the gene level, can take longer to be effective than reagents that target at the transcript level; further, genetic mutations induced by CRISPR-Cas9, though effective at knocking out gene expression and/or function, can induce genetic compensation for some gene targets, which typically does not occur with knockdown at the RNA or protein level (Rossi et al., 2015). While preexisting RNA-targeting tools, most notably RNA interference (RNAi), have been shown to be effective in other model organisms, such as roundworm (*Caenorhabditis elegans*), fruit fly (*Drosophila melanogaster*) and mouse (*Mus musculus*) (Perrimon et al., 2010), it has had little success in the chick (Hernandez and Bueno, 2005). Thus, there is a need among avian researchers for a plasmid-based, alternative knockdown approach that functions at the transcript level to elicit effective gene expression knockdown.

Cas13, a class 2 type VI CRISPR-Cas RNA endonuclease, functions by forming a ribonucleoprotein complex with a single guide RNA (gRNA) to cleave target RNAs which, when targeted to the coding region, can disrupt translation and protein production (Wessels et al., 2020). Prior studies have successfully implemented Cas13 as a reliable method to knock down gene expression in mammalian cell lines, demonstrating higher efficacy and specificity compared to RNAi (Abudayyeh et al., 2017; Cox et al., 2017; Konermann et al., 2018). Recently, Cas13d, a subtype of the Cas13 family, has been successfully adapted for use in intact animal models, most notability in zebrafish (*Danio rerio*) embryos, to knock down gene function with specificity while avoiding embryonic toxicity. Cas13d is especially valuable as a loss-of-function genetic tool in zebrafish, for which RNAi has failed to become an effective knockdown method. This novel technology, initially adapted for zebrafish, has also been demonstrated to be compatible with other animal models such as medaka (*Oryzias latipes*), killifish (*Nothobranchius furzeri*), and mouse (*Mus musculus*) embryos (Kushawah et al., 2020).

Despite its applicability across a broad range of animal models, no prior study has explored the potential of CRISPR-Cas13d in an avian model. Here, we present our adaptation of the CRISPR-Cas13d system to achieve targeted gene expression knockdown in chick embryos. As proof-of-principle, we demonstrate the knockdown of PAX7, an early neural crest cell marker. We designed and implemented a two-plasmid delivery approach, co-electroporating a Cas13d plasmid and a guide RNA plasmid to express the knockdown machinery in the chick. Expression of our CRISPR-Cas13d system *in vivo* resulted in significant knockdown of PAX7 expression and function, as evidenced by a reduction of neural crest migration away from the midline, which is a functional consequence of the loss of PAX7 (Basch et al., 2006). CRISPR-Cas13d complements preexisting knockdown tools such as CRISPR-Cas9 and morpholino oligonucleotides, thereby expanding the experimental potential and versatility of the avian model system.

## 2. MATERIALS AND METHODS

### 2.1 Guide RNA design

The guide RNA (gRNA) design protocol was adapted from previously published methods (Hernandez-Huertas et al., 2022; Kushawah et al., 2020). The full-length *Pax7* mRNA and coding sequences (CDS) for *Gallus gallus* were obtained from the National Center for Biotechnology Information (NCBI; Accession: NM_205065.1). RNAfold [http://rna.tbi.univie.ac.at/cgi-bin/RNAWebSuite/RNAfold.cgi; (Lorenz et al., 2011)] was first used to predict secondary structure and the accessible regions of full-length *Pax7* mRNA. Candidate *Pax7* gRNA sequences targeting the coding region were then generated using cas13design (https://cas13design.nygenome.org). Each candidate gRNA from cas13design was screened against the predicted secondary structure from RNAfold and three unique gRNAs that best targeted accessible regions across the *Pax7* coding region were selected. The Control gRNA sequence was taken from standard control morpholino (MO) sequence manufactured by Gene Tools, which is predicted to have no sequence complementarity in chick.

### 2.2 Molecular cloning

To generate a donor plasmid, pCAG-memRFP (**Fig. 2A**), we digested pCI-H2B-RFP (Betancur et al., 2010) with NheI and NotI restriction enzymes to excise the internal ribosome entry site (IRES) and H2B-RFP coding region. We then inserted a fragment encoding a membrane-localized RFP (memRFP) and a multiple cloning site (MCS), which included AgeI, ClaI, HindIII, and NotI restriction sites. The donor plasmid was then digested with a combination of AgeI/ClaI or NheI/NotI restriction enzymes to supply the vector backbone required for generating the gRNA (**Fig. 2B-C**) and Cas13d plasmids (**Fig. 2C-D**), respectively. All inserts were commercially synthesized by Twist Biosciences as clonal genes or gene fragments; gene fragments were directly cloned into pCR™-Blunt II-TOPO™ vector for amplification and restriction digestion, except for the gRNA3 insert, which was PCR amplified prior to restriction digest.

**Figure 1.**
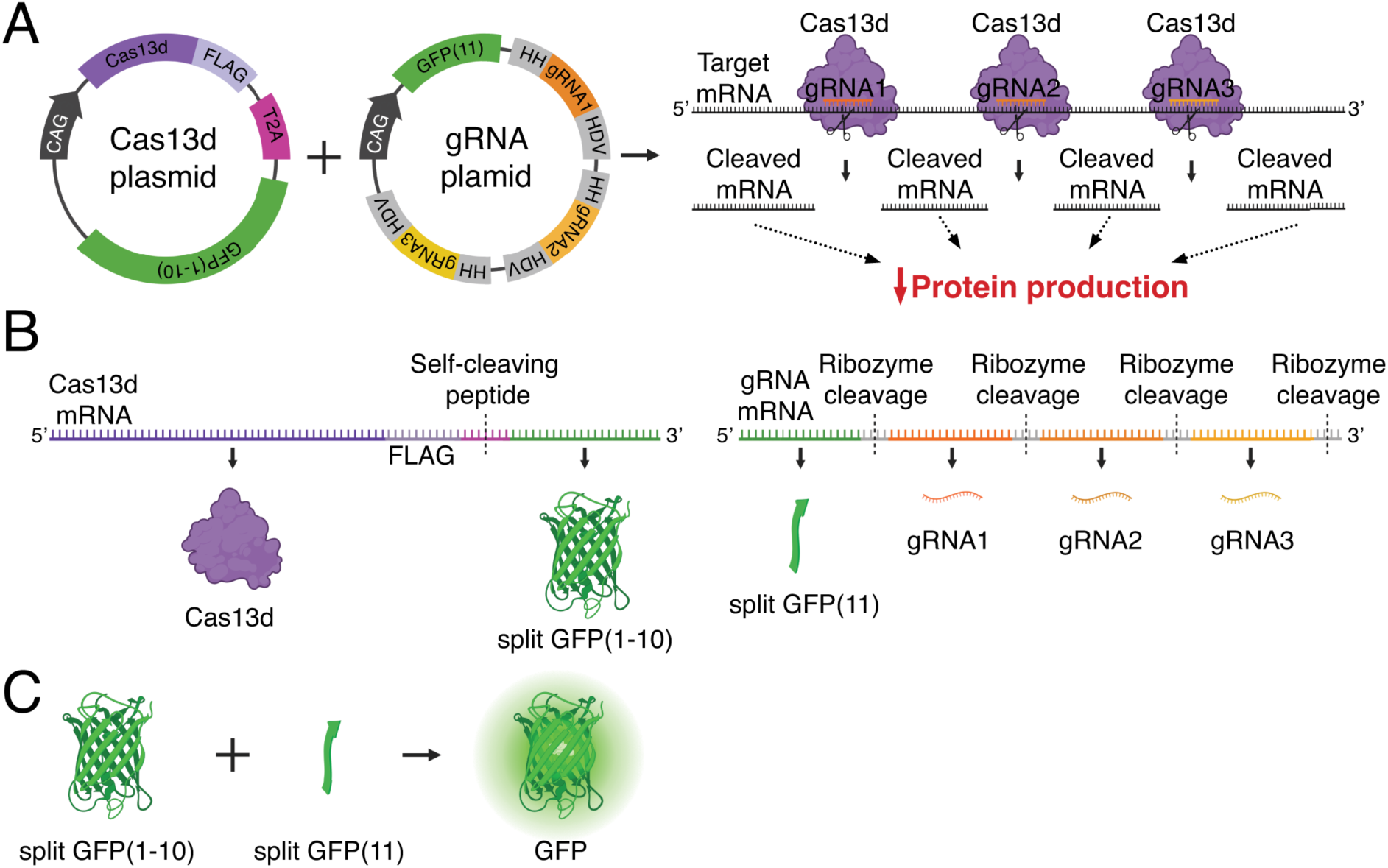
Delivery strategy of a two-plasmid CRISPR-Cas13d system for avian embryos. (**A**) Schematic depicting structure of Cas13d and guide RNA (gRNA) plasmids and the *in vivo* knockdown effect. Plasmid expression is driven by a ubiquitous promoter (CAG). Cas13d protein and gRNAs form ribonucleoprotein complexes to cleave target mRNA at multiple sites across the coding region, ultimately resulting in disrupted translation and decreased protein production. (**B**) The Cas13d plasmid generates a FLAG-tagged Cas13d protein and a split GFP(1-10) non-fluorescent reporter protein, separated by a T2A self-cleaving peptide. The gRNA plasmid generates a split GFP(11) non-fluorescent reporter protein, and three unique gRNAs flanked by ribozyme sequences (HH and HDV) that separate the transcribed components. (**C**) When co-expressed, the split GFP non-fluorescent reporters GFP(1-10) and GFP(11) self-complement into a functional, fluorescent GFP reporter protein to label cells that have successfully received both plasmids. HH, hammerhead ribozyme sequence; HDV, hepatitis delta virus ribozyme sequence.

**Figure 2.**
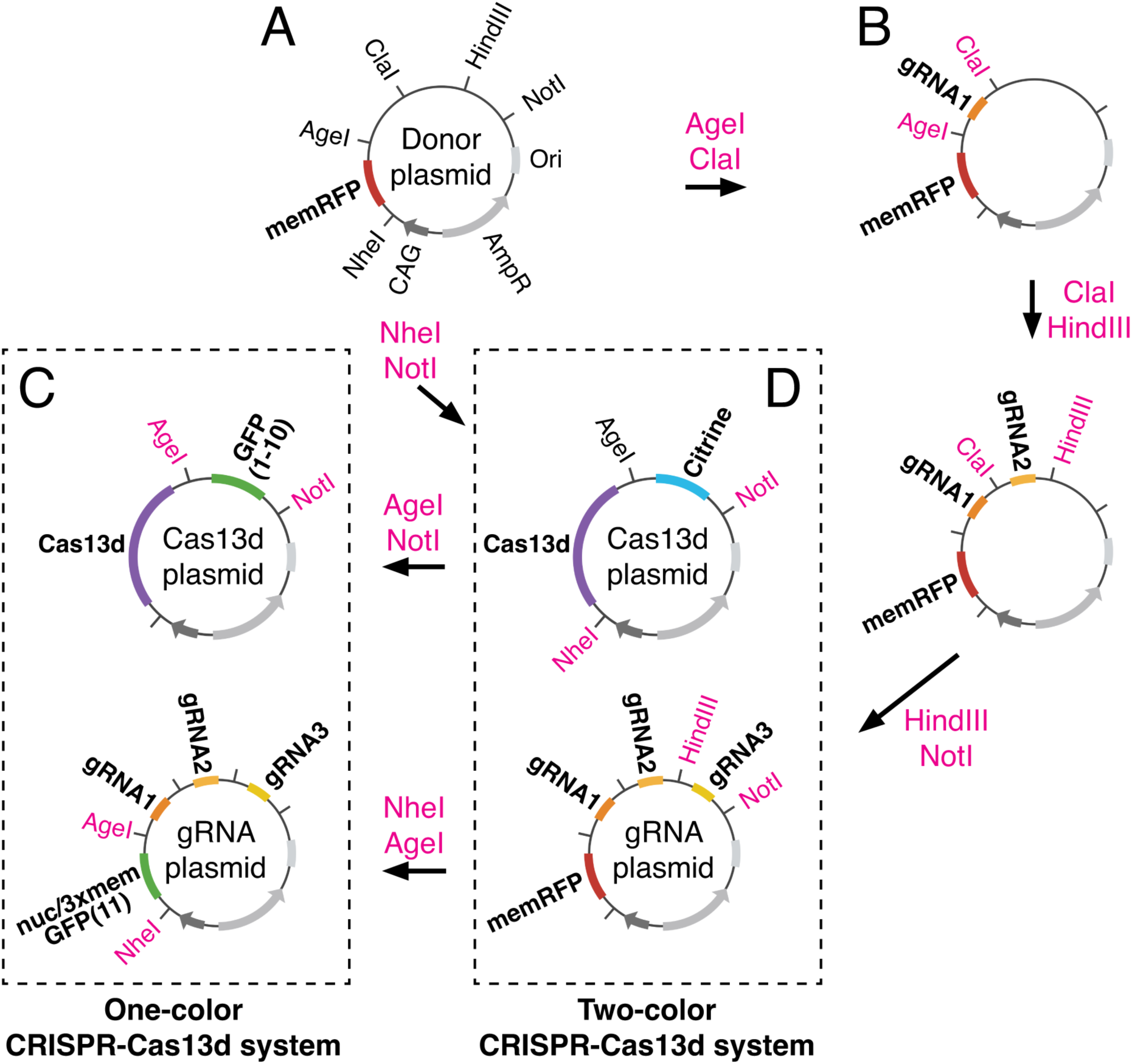
Sequential cloning strategy used to generate one- and two-color CRISPR-Cas13d systems. (**A**) Donor plasmid supplying the backbone vector contains the following features: Origin of replication (Ori), Ampicillin resistance (AmpR), ubiquitous promoter (CAG), and a membrane-localized RFP reporter (memRFP), as well as a multiple cloning site. Relevant restriction sites are shown. (**B-D**) Donor plasmid was digested with indicated restriction enzymes and ligated iteratively to insert three guide RNAs (gRNAs). Three variations of the gRNA plasmid were created: one containing memRFP for a two-color system (**D**), and two others containing split GFP(11) reporters that are either nuclear (nucGFP(11)) or membrane localized (3xmemGFP(11)) for a one-color system (**C**). Donor plasmid was also separately digested with indicated restriction enzymes and ligated with respective inserts to create two variations of the Cas13d plasmid: one containing a split GFP reporter (GFP(1-10)) for the one-color system (**C**), and the other containing a Citrine reporter for the two-color system (**D**).

The Cas13d-FLAG-T2A-Citrine insert with NheI/NotI restriction sites was purchased as a custom clonal gene plasmid, then cloned into the digested donor plasmid (**Fig. 2D**). We then exchanged the Citrine coding region for split GFP(1-10) via AgeI and NotI restriction sites to generate the pCAG-Cas13d-T2A-GFP(1-10) plasmid (**Fig. 2C**). To generate gRNA constructs, we commercially synthesized individual fragments that contained the gRNA sequence (constant direct repeat stem loop sequence followed by a variable spacer sequence complementary to the target) flanked by hammerhead (HH) and hepatitis delta virus (HDV) ribozyme self-cleavage sites and restriction sites for directional cloning on either side. We then sequentially cloned each gRNA into the donor plasmid to generate a gRNA plasmid containing coding for a fluorescent protein reporter (memRFP) followed by three tandem gRNAs that would be separated from the mRNA via ribozyme cleavage after transcription (**Fig. 2B, D**). We then exchanged the memRFP coding region for three tandem split GFP(11) proteins, containing either a membrane localization signal or histone H2B (for nuclear localization) via NheI and AgeI restriction sites (**Fig. 2C**). Combination of the Citrine-or split GFP(1-10)-containing Cas13d plasmid with the memRFP-or split GFP(11)-containing gRNA plasmid generates a two-color (**Fig. 2D**) or single-color CRISPR-Cas13d system (**Fig. 2C**), respectively.

### 2.3 Electroporation

Fertile chicken eggs (*Gallus gallus*) were purchased locally (Petaluma Farms, Petaluma, CA). Prior to *ex ovo* electroporation, eggs were incubated in a humidified 100°F incubator and electroporated at stage HH4 by passing 5.0 V pulses for 50 ms each every 100 ms, using previously described techniques (Sauka-Spengler and Barembaum, 2008). For Cas13d-mediated PAX7 knockdown, Cas13d plasmid (pCAG-Cas13d-T2A-GFP(1-10)) [1 μg/μL] was co-electroporated with either a Control gRNA plasmid (pCAG-nucGFP11-3xControlgRNA) [4.5 μg/μL] on the left side of the embryo, or a Pax7 gRNA plasmid (pCAG-nucGFP11-3xPax7gRNA) [4.5 μg/μL] on the right side of the embryo. Embryos were then screened following incubation for nuclear GFP fluorescence, which indicated electroporated cells received both Cas13d and gRNA plasmids. For morpholino (MO)-mediated PAX7 knockdown, the left side of the embryo was co-electroporated with [1.2 mM] standard control MO (Gene Tools) and 2 μg/μL pCI-H2B-RFP, while the right side of the embryo was co-electroporated with [1.2 mM] Pax7 MO [Gene Tools; (Basch et al., 2006; Roellig et al., 2017)] and 2 μg/μL pCIG (Megason and McMahon, 2002). Electroporated embryos were cultured in 1 mL of albumin supplemented with 1% penicillin-streptomycin in an incubator set to 99°F. The incubator was turned on for the first 2 hours immediately following electroporation to initiate *in vivo* synthesis of CRISPR components. The incubator was then turned off for 8 hours to allow sufficient time for the expression and activation of CRISPR machinery. Finally, the incubator was turned back on for an additional 8 hours, resulting in a total incubation time of ∼10 hours post-electroporation, or until the embryos reached stage HH9/9+ according to Hamburger–Hamilton staging (Hamburger and Hamilton, 1951).

### 2.4 Immunostaining

For whole mount immunostaining, embryos were fixed at room temperature for 20 min with 4% paraformaldehyde in 0.1M sodium phosphate buffer (PFA). Washes, blocks (10% donkey serum), and antibody incubations were performed with TBSTx (0.5M Tris-HCl/1.5M NaCl/10mM CaCl_2_/0.5% Triton X-100/0.001% Thimerosal) as previously described (Chacon and Rogers, 2019; Manohar et al., 2020). For cross-sections, immunostained embryos were post-fixed in PFA at 4°C overnight, and washed, embedded, and cryo-sectioned as previously described (Hutchins and Bronner, 2018, 2019). Specific antibodies and reagents are indicated in the Key Resources Table.

### 2.5. Image acquisition

Epifluorescence images were acquired using a Zeiss Axio Imager M2 with an Apotome 3 module. Whole mount embryos were imaged with a 10x objective lens (Zeiss Plan-Apochromat 10x, NA=0.45), and transverse cross-sections were imaged with a 20x objective lens (Zeiss Plan-Apochromat 20x, NA=0.8). Apotome-processed Z-stacks of whole mount embryos and cross-sections were converted into maximum intensity projections for display. Images were minimally processed for brightness/contrast and pseudo-colored using Fiji (ImageJ, NIH) (Schindelin et al., 2012) and Adobe Photoshop CC.

### 2.6 Quantification and statistical analysis

All quantifications shown in this study were performed on maximum intensity projections in Fiji. Statistical analyses were performed in Prism 10 (GraphPad). To quantify PAX7 knockdown in transverse cross-sections as shown in Figure 3B, we used the freehand selection tool to outline regions of interest around PAX7^+^ cells and measured the integrated density of PAX7 fluorescence within those regions. The PAX7 intensity measurements of each embryo (n = 15) are averages of the integrated densities obtained from three nonadjacent cross-sections per embryo. Relative reduction in PAX7 intensity was calculated by dividing the average PAX7 intensity detected on the right side (PAX7 knockdown) by the left side (contralateral control) of individual embryos. One-sample Wilcoxon signed-rank test was performed to determine statistical significance.

**Figure 3.**
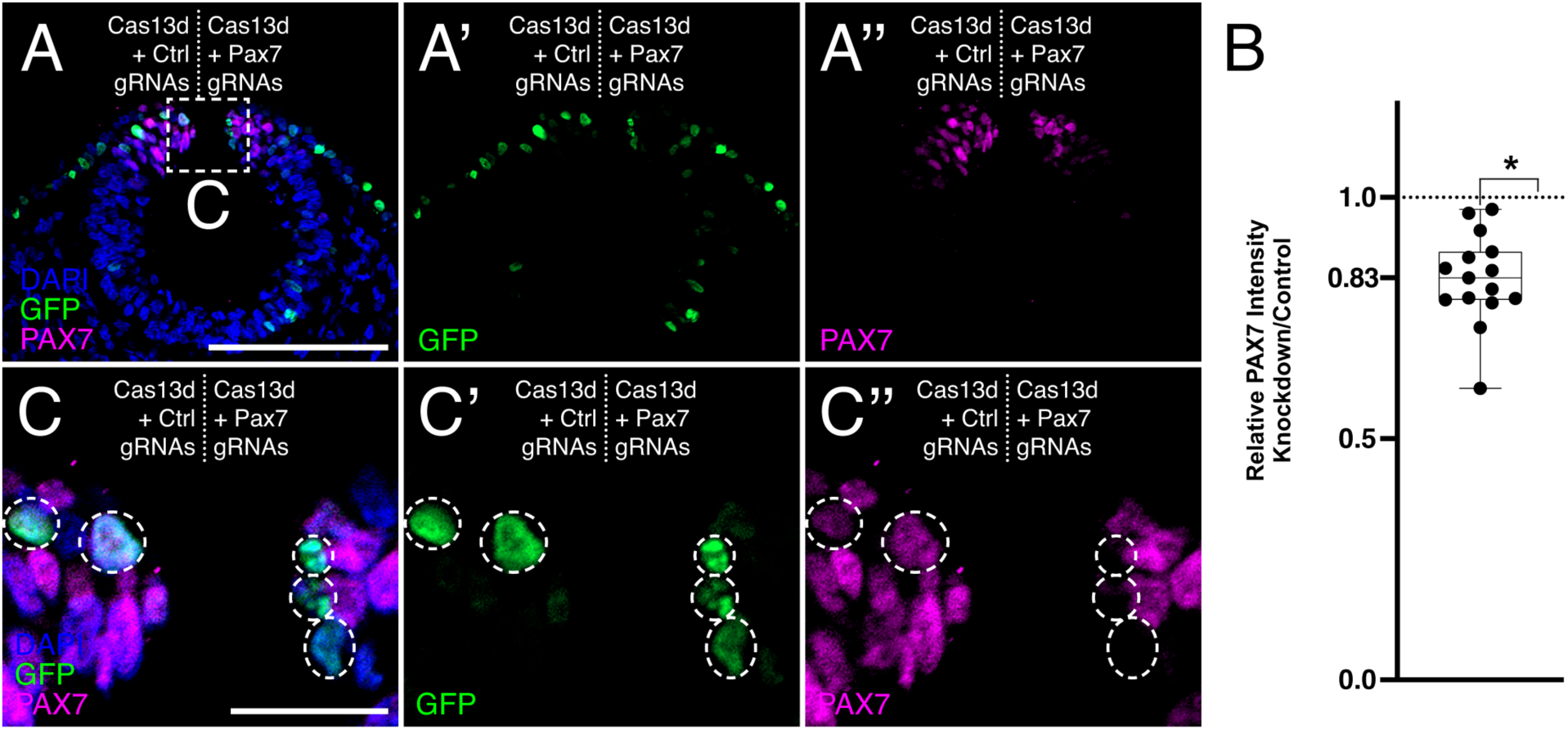
One-color CRISPR-Cas13d-mediated PAX7 knockdown. (**A**) A representative transverse cross-section of a HH9 chick embryo head bilaterally electroporated with Cas13d + Control guide RNA (gRNA) plasmids (left), and Cas13d + Pax7 gRNA plasmids (right). Reduced PAX7 intensity is observed on the right side of the embryo. Scale bar, 100 μm. (**B**) Quantification of PAX7 knockdown. Each data point represents the relative PAX7 intensity detected on the right side of an embryo (PAX7 knockdown) compared to the left side of the same embryo (contralateral control). The relative intensity measurements for each embryo are averages of the intensities detected from three nonadjacent cross-sections. A statistically significant reduction in PAX7 intensity is observed (83.1 ± 2.4% of the contralateral control side; n=15 embryos) with CRISPR-Cas13d-mediated knockdown. *, p < 0.0001; one-sample Wilcoxon signed-rank test. (**C**) Enlarged view of boxed area shown in (**A**). GFP^+^ cells are highlighted by white circles. Only the (**C’**) GFP^+^ cells on the right side of the embryo show reduction in (**C’’**) PAX7 intensity. Scale bar, 20 μm.

To quantify functional knockdown of PAX7 in whole mount embryos as shown in Figure 4E, we used the wand tool (set to 8-connected mode) to trace PAX7^+^ area, which represents the area of neural crest migration. Relative reduction in this area was calculated by dividing the PAX7^+^ area detected on the right side (PAX7 knockdown) by the left side (contralateral control) of individual embryos. One-sample Wilcoxon signed-rank test was performed to determine statistical significance. Mann-Whitney test was performed to compare the degree of functional knockdown achieved with MO (n = 11) versus CRISPR-Cas13d (n = 13).

**Figure 4.**
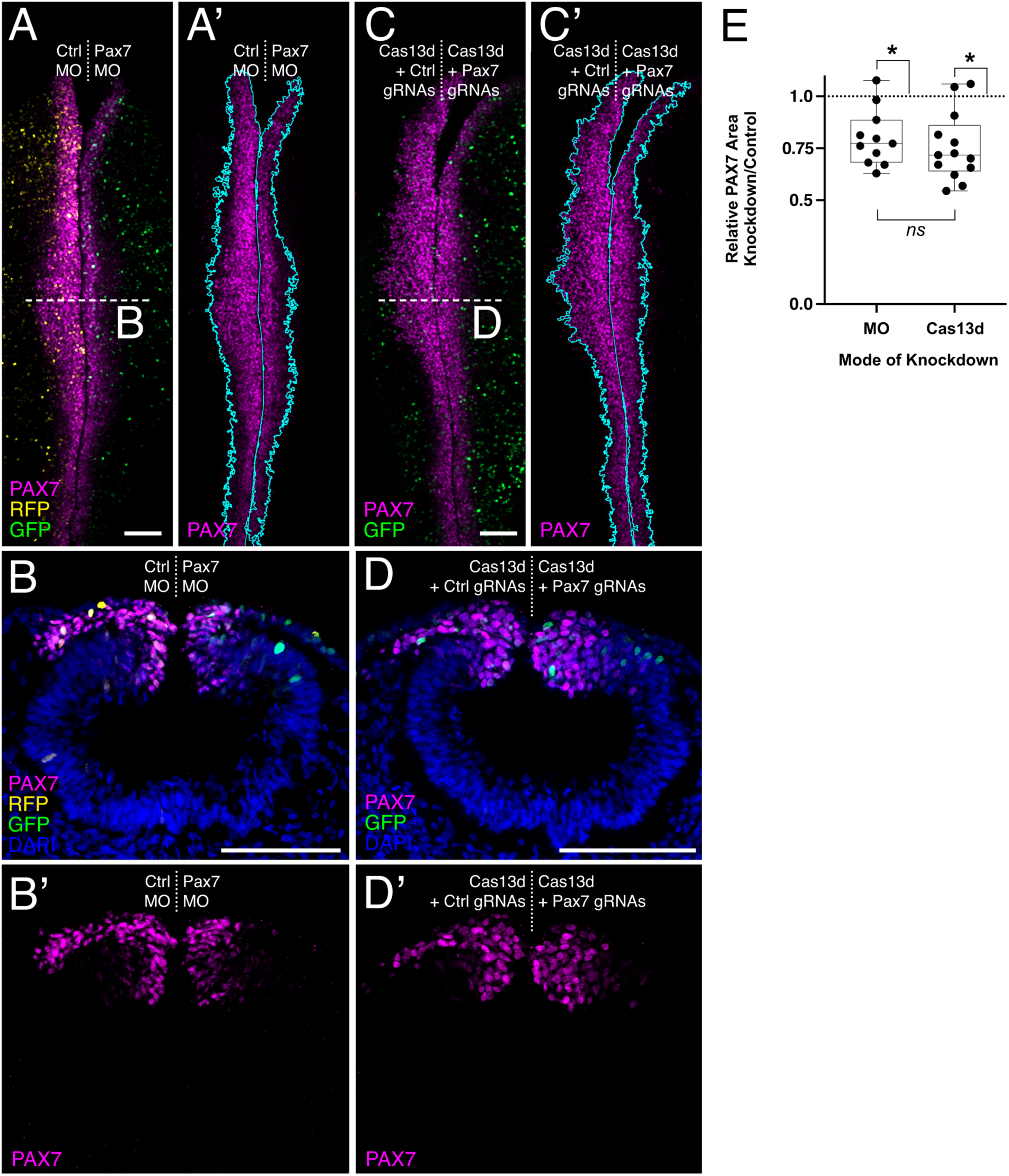
A comparison of morpholino- and CRISPR-Cas13d-mediated PAX7 knockdown approaches. (**A-D**) Representative maximum intensity projections of HH9+ chick embryos electroporated with morpholino (MO) or CRISPR-Cas13d reagents and immunostained for PAX7 (magenta). (**A**) MO knockdown embryos were bilaterally electroporated with a control morpholino (Ctrl MO) and an RFP reporter (left), and Pax7 MO and a GFP reporter (right). (**B**) Representative transverse cross-section of embryo shown in (**A**). (**C**) One-color CRISPR-Cas13d knockdown embryos were electroporated with Cas13d + Control gRNA plasmids (left), and Cas13d + Pax7 gRNA plasmids (right), using split GFP fluorescence to indicate co-expression of both plasmids. Cyan outlines the area of PAX7^+^ cells, which is used to calculate the area of neural crest migration. (**D**) Representative transverse cross-section of embryo shown in (**C**). (**E**) Quantification of the neural crest migration defect. Each data point represents the relative area of neural crest migration away from the midline, indicated by PAX7^+^ cells, detected on the right side of an embryo (PAX7 knockdown) compared to the left side of the same embryo (contralateral control). A statistically significant reduction in PAX7^+^ migration area is observed with MO-mediated knockdown of PAX7 (80.0 ± 4.1% of PAX7^+^ cell migration area compared to control; n=11 embryos). *, p = 0.0029, one-sample Wilcoxon signed-rank test. Similarly, a statistically significant reduction in PAX7^+^ migration area is also observed with CRISPR-Cas13d-mediated knockdown of PAX7 (75.5 ± 4.5% of PAX7^+^ cell migration area compared to control; n=13 embryos). *, p = 0.0012, one-sample Wilcoxon signed-rank test. Notably, the defects observed in neural crest migration for the two modes of knockdown are not different. *ns*, nonsignificant; p = 0.2066, Mann-Whitney test. Scale bar, 100 μm (**A, C**); 20 μm (**B, D**).

### 2.7 Plasmid availability

Donor, Cas13d, and Control and Pax7 guide RNA (gRNA) plasmids for the one- and two-color CRISPR-Cas13d systems will be made available through Addgene (https://www.addgene.org) upon publication. The catalog numbers for the plasmids described in this study can be found in the Key Resources Table.

## 3. RESULTS AND DISCUSSION

### 3.1 Two-plasmid delivery approach

Our adapted CRISPR-Cas13d system is based on a recently developed CRISPR approach that applied direct injection of *in vitro* synthesized components to knock down gene expression in fish and mouse embryos (Kushawah et al., 2020). To extend the use of this CRISPR system to avian embryos, we have modified this system and here introduce a two-plasmid delivery approach to induce efficient gene expression knockdown. Like the previously optimized CRISPR-Cas9 system (Gandhi et al., 2017), our novel two-plasmid CRISPR-Cas13d system consists of a Cas13d-expressing plasmid and a guide RNA (gRNA)-expressing plasmid, which are co-electroporated into the chick embryo. Once electroporated, the CAG promoter (chicken beta-actin promoter and CMV IE enhancer) (Hitoshi et al., 1991; Sauka-Spengler and Barembaum, 2008) drives robust, ubiquitous expression of the CRISPR-Cas13d elements: 1) Cas13d protein, an RNA endonuclease that complexes with a single, short, sequence-specific gRNA to target and cleave an mRNA transcript; and 2) three unique gRNAs that are complementary to multiple sites with the coding region of a single target. When co-expressed and complexed together, Cas13d is targeted to and cleaves the mRNA, impeding translation and effectively decreasing protein expression (**Fig. 1A**).

An additional aspect of our CRISPR-Cas13d system is the use of one-or two-color fluorescent reporter proteins to visually identify cells that received both plasmids. For the one-color reporter system, we developed a Cas13d plasmid that produces Cas13d protein and split GFP(1-10), a non-fluorescent split GFP protein containing the first ten GFP β-strands, separated from Cas13d via the T2A self-cleaving peptide sequence (Gandhi et al., 2021; Williams et al., 2018). The gRNA plasmid supplies a gRNA transcript that encodes the remaining eleventh GFP β-strand, a non-fluorescent nuclear-localized split GFP(11) protein (nucGFP(11)), and three unique gRNAs in tandem. Each gRNA is flanked by hammerhead (HH) ribozyme and hepatitis delta virus (HDV) ribozyme sequences on the 5’ and 3’ ends, respectively (**Fig. 1B**); the precursor gRNA transcript undergoes self-catalyzed ribozyme cleavage to release three individual, functional gRNAs (Gandhi et al., 2021; He et al., 2017). When expressed in the same cells, the GFP(1-10) from the Cas13d plasmid and the nucGFP(11) from the gRNA plasmid self-complement to form a stable, fluorescent GFP reporter that localizes to the nucleus (**Fig. 1C**; (Feng et al., 2017)). To validate the split GFP reporter *in vivo* and ensure fluorescence is produced only when Cas13d and gRNA plasmids are co-expressed, we performed bilateral electroporations with the Cas13d plasmid alone on the left side of the embryo and the combined Cas13d and gRNA plasmids on the right side of the embryo; we found that endogenous GFP fluorescence is only detectable when both the Cas13d and gRNA plasmids are present, whereas immunostaining in another fluorescence channel with an antibody against GFP that recognizes GFP(1-10) detects the presence of the Cas13d plasmid with or without the gRNA plasmid (**Supplemental Fig. S1**). Thus, the use of split GFP self-complementation allows detection of CRISPR-Cas13d knockdown reagents in a single fluorescence channel.

We also designed a two-color CRISPR-Cas13d system, which expresses Citrine and membrane-localized RFP (memRFP) from the Cas13d and gRNA plasmids, respectively (**Supplemental Fig. S2**). The two-color system functions similarly to the one-color system, and offers researchers versatility based on their experimental needs.

### 3.2 Cloning strategy

The combined Cas13d and gRNA plasmids encode the knockdown machinery required for a fully functional CRISPR-Cas13d system. We engineered these plasmids to be modular, creating a donor plasmid that contains the CAG promoter, a membrane-localized RFP reporter (memRFP), and a multiple cloning site to allow for restriction enzyme-based directional cloning (**Fig. 2A**). We first generated our gRNA plasmid using a sequential cloning strategy with the donor plasmid to insert gRNA fragments (containing the flanking ribozyme sequences) in the correct orientation (**Fig. 2B**). Due to the repetitive ribozyme sequences, we were unable to commercially synthesize a single insert containing multiple gRNA sequences, and thus found this to be the most straightforward and inexpensive solution to gRNA plasmid synthesis. The modular gRNA plasmid construction also allows researchers to exchange gRNA sequences with relative ease (**Fig. 2C-D**).

Starting from the donor plasmid, we also generated two variations of the Cas13d plasmid for use in the one-color or two-color system (**Fig. 1; Fig. 2C-D; Supplemental Fig. S2**). We engineered a Cas13d plasmid containing a Citrine reporter for use with the memRFP-containing gRNA plasmid in the two-color system, inserting a fragment encoding Cas13d-FLAG-T2A-Citrine in place of memRFP in the donor plasmid. We then modified this Cas13d plasmid by exchanging the Citrine for the split GFP(1-10) via AgeI and NotI restriction sites for use in the one-color system; we designed these Cas13d plasmids to have the AgeI site 3’ of the T2A site and remain in-frame, so that researchers can easily exchange reporter elements.

In a similar manner, three variations of the gRNA plasmid were also generated: one containing a memRFP reporter (as described above; **Fig. 2B**), and two others containing variations of split GFP(11) reporters—one that is nuclear localized (nucGFP(11)) and one that is membrane localized (3xmemGFP(11)) (**Fig. 2C-D**). These variations of the split GFP reporter result in different subcellular localization of self-complemented GFP *in vivo*, which could allow researchers the versatility to perform knockdowns for more than one target, or increase the number of gRNAs for a single target, while maintaining the ability to distinguish cells co-expressing multiple gRNA plasmids. These examples showcase just a few of the many ways the CRISPR-Cas13d system can be adapted to meet a variety of experimental needs in the avian model system.

### 3.3 PAX7 knockdown

As proof-of-principle, we used our two-plasmid CRISPR-Cas13d system in the chick embryo to demonstrate knockdown of PAX7, an early neural crest cell marker (**Fig. 3; Supplemental Fig. S2**). We designed three unique gRNAs targeting the coding sequence of *Pax7* mRNA and generated gRNA plasmids using the cloning strategy described in (**Fig. 2**). We delivered the one-color CRISPR-Cas13d system to the chick embryo via bilateral *ex ovo* electroporation at Hamburger–Hamilton stage 4 (HH4). Since electroporated constructs require time to be transcribed and translated into functional CRISPR components, we additionally implemented a developmental “pause” post-electroporation by modulating incubation time and temperature. This optimized incubation strategy allows sufficient time for expression and activation of the CRISPR machinery. Additionally, the chick embryo can be bilaterally electroporated with different gRNA constructs, allowing for an internal control within a single embryo. Here, the right side of the chick embryo was targeted for PAX7 knockdown and was co-electroporated with Cas13d and Pax7 gRNA split GFP reporter plasmids, whereas the contralateral left side was co-electroporated with Cas13d and Control gRNA split GFP reporter plasmids.

We examined electroporated embryos at HH9/9+ via immunostaining for PAX7 and first compared the relative fluorescence intensity of PAX7 staining between the right (knockdown) and left (control) sides of the embryo. In transverse cross-sections, the right side of chick embryo showed reduced PAX7 fluorescence intensity in the dorsal neural tube at HH9 (83.1 ± 2.4% of the contralateral control side) (**Fig. 3A-3A’’**). This ∼17% reduction in PAX7 expression was statistically significant (p < 0.0001, n=15 embryos) (**Fig. 3B**) and comparable to the reduction achieved in previous work using translation-blocking morpholino targeting *Pax7* (Roellig et al., 2017). Further, GFP^+^ cells successfully co-electroporated with Cas13d and Pax7 gRNA plasmids, as indicated by GFP fluorescence from self-complementation, show drastic reduction in PAX7 levels relative to surrounding GFP^—^ cells, whereas cells co-electroporated with the Control gRNA plasmid do not (**Fig. 3C-3C’’**), further demonstrating the specificity of our CRISPR-Cas13d system.

Given the mosaic nature of electroporations, it is important to note that the quantification of PAX7 knockdown described in (**Fig. 3B**) is an underestimation of knockdown efficiency, as it includes cells that did not necessarily receive both the Cas13d and gRNA plasmid, to assess knockdown as conservatively as possible. For this reason, we also assessed the functional knockdown of PAX7. PAX7 is an early marker of neural crest cells, and its reduction using Pax7 MO leads to a decrease in neural crest migration (Basch et al., 2006). To assess the severity of functional defect resulting from CRISPR-Cas13d-mediated PAX7 knockdown, we targeted PAX7 separately with MO or CRISPR-Cas13d and compared neural crest migration deficits, as indicated by the area of PAX7^+^ cell migration away from the midline at HH9+. As described above with CRISPR-Cas13d-mediated knockdown, we performed bilateral *ex ovo* electroporation at stage HH4 with translation-blocking Pax7 MO (Basch et al., 2006; Roellig et al., 2017) and standard control MO. The right side of the chick embryo was co-electroporated with Pax7 MO and pCIG, which encodes a nuclear GFP reporter (Megason and McMahon, 2002). Its contralateral left side was co-electroporated with standard control MO and pCI-H2B-RFP, which encodes a nuclear RFP reporter (Betancur et al., 2010). As with CRISPR-Cas13d, MO-electroporated embryos were similarly harvested and immunostained for PAX7.

In whole mount embryos, both modes of knockdown showed decreased neural crest migration relative to their contralateral control sides, as indicated by the area of PAX7^+^ cell migration away from the midline (**Fig. 4A-E**). MO-mediated knockdown yielded a ∼20% reduction (80.0 ± 4.1% of PAX7^+^ cell migration area compared to control; p = 0.0029, n=11 embryos) and CRISPR-Cas13d-mediated knockdown exhibited a ∼25% reduction (75.5 ± 4.5% of PAX7^+^ cell migration area compared to control; p = 0.0012, n=13 embryos) in neural crest cell migration area. Notably, the severity of functional defect resulting from PAX7 knockdown was not significantly different between the two methods of knockdown (p = 0.2066), demonstrating the efficacy and utility of our CRISPR-Cas13d system. Thus, we have successfully designed and implemented a two-plasmid CRISPR-Cas13d system for use in the avian embryo to knockdown gene expression.

## 4. CONCLUSIONS

In this study, we present a novel method of gene expression knockdown optimized for use in avian embryos, the CRISPR-Cas13d system. The chick embryo is a useful model system for functional gene analysis due its external development, which allows experimental perturbations at early developmental stages, and homology to human development. Capitalizing on the unique advantages of the chick embryo, we designed a two-plasmid CRISPR-Cas13d system for efficient gene expression knockdown *in vivo*. Given that the *in vivo* expression of CRISPR components is driven by the chick’s endogenous transcriptional machinery, future applications utilizing this method could potentially impose knockdown in a tissue-specific or drug-inducible manner, via the control of various enhancers or alternative promoters. In summary, we demonstrate that our adaptation of the CRISPR-Cas13d system is a useful method for achieving efficient and specific gene expression knockdown in the avian embryo. This alternative mode of knockdown complements preexisting loss-of-function genetic tools, such as CRISPR-Cas9 and morpholinos, thereby expanding the experimental potential and versatility of the avian model system.

## Supporting information

Key Resources Table

## CRediT AUTHORSHIP CONTRIBUTION STATEMENT

**Minyoung Kim:** Writing – original draft, Visualization, Methodology, Investigation, Formal analysis, Data curation. **Erica J. Hutchins:** Writing – review & editing, Supervision, Resources, Methodology, Investigation, Funding acquisition, Conceptualization.

## ACKNOWLEDGMENTS

The authors are supported by the National Institutes of Health (NIH) R00DE028592 (EJH), R35GM150763 (EJH), and Institutional Training Grant 5T32DE007306 (MK). We thank Dr. Michael Piacentino for critical input. Schematics were created with BioRender.com.

---

**Supplemental Figure S1.**
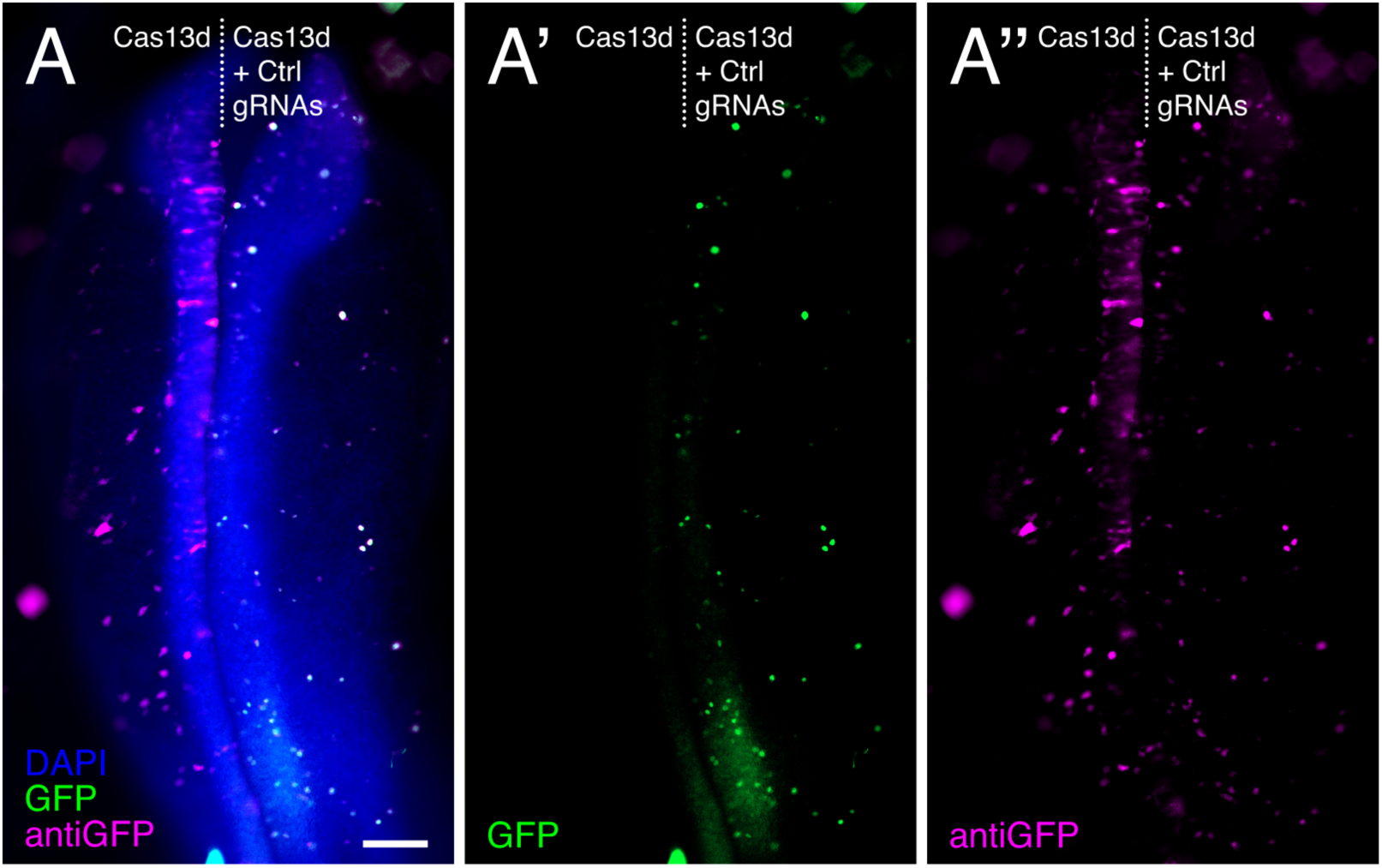
Validation of split GFP reporter system. (**A-A’’**) A representative epifluorescence micrograph of a whole mount HH9 chick embryo head bilaterally electroporated with the one-color CRISPR-Cas13d system using split GFP, using Cas13d plasmid alone (left) and Cas13d + Control gRNA plasmids (right). Fluorescent GFP signal is detected only on the right side of the embryo for n=2/2 embryos. Scale bar, 100 μm.

**Supplemental Figure S2.**
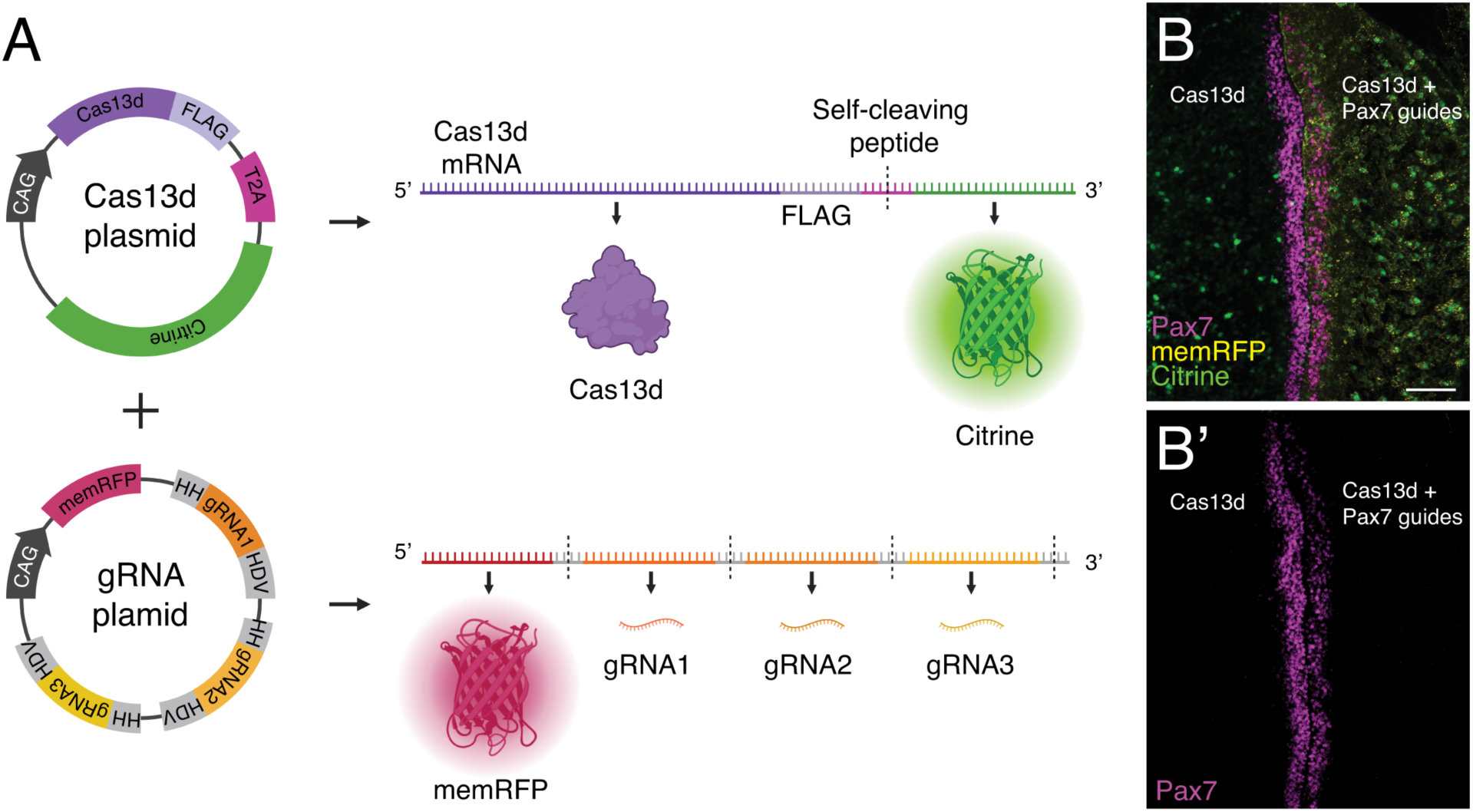
Delivery strategy of the two-color CRISPR-Cas13d system. (**A**) Schematic depicting structure of Cas13d and guide RNA (gRNA) plasmids for the two-color system. Plasmid expression is driven by a ubiquitous promoter (CAG). The Cas13d plasmid generates a FLAG-tagged Cas13d protein and a Citrine fluorescent reporter protein, separated by a T2A self-cleaving peptide. The gRNA plasmid generates a membrane-localized RFP fluorescent reporter protein, and three unique gRNAs flanked by ribozyme sequences (HH and HDV) that separate the transcribed components. (**B**) Embryo (HH9-) bilaterally electroporated with the Cas13d plasmid on both sides while the right-side additionally received the Pax7 gRNA plasmid. PAX7 knockdown is indicated by reduced PAX7 staining (**B’**) on the right. HH, hammerhead ribozyme sequence; HDV, hepatitis delta virus ribozyme sequence. Scale bar, 100 μm.

