## Supplementary material for "CRISPR-Cas13d as a molecular tool to achieve targeted gene expression knockdown in chick embryos": Key Resources Table

| Reagent or resource | Source | Identifier |
| --- | --- | --- |
| Antibodies | | |
| Mouse IgG1 anti-PAX7 | Developmental Studies Hybridoma Bank | Cat#: pax7  RRID: AB_528428 |
| Goat IgG anti-GFP | Rockland | Cat#: 600-101-215  RRID: AB_218182 |
| Rabbit IgG anti-FLAG | Millipore/Sigma | Cat#: F7425  RRID: AB_439687 |
| Donkey anti-Goat IgG Alexa Fluor 647 | Invitrogen | Cat#: A21447 |
| Goat anti-Mouse IgG1 Alexa Fluor 647 | Invitrogen | Cat#: A21240 |
| Goat anti-Rabbit IgG Alexa Fluor 568 | Invitrogen | Cat#: A11011 |
| Chemicals, Peptides, and Recombinant Proteins | | |
| DAPI | Thermo Fisher | Cat#: D1306 |
| Fluoromount-G | SouthernBiotech | Cat#: 0100-01 |
| Experimental Models: Organisms/Strains | | |
| *Gallus gallus* | Petaluma Farms (Petaluma, CA) | Fertile Rhode Island Red eggs |
| Oligonucleotides | | |
| Morpholino: Control  5’-CCTCTTACCTCAGTTACAATTTATA | GeneTools |  |
| Morpholino: Pax7  5’-TCCGTGCGGAGCGGGTCACCCCC | GeneTools; Basch et al., 2006 |  |
| Primer: gRNA3  Forward: 5’-GTAATAGCGAAGAGGCCCGC  Reverse: 5’-ATAGCGGCCGCTCATCTAGA | This paper; IDT | Custom DNA Oligos |
| Recombinant DNA | | |
| Plasmid: pCI-H2B-RFP | Betancur et al., 2010 |  |
| Plasmid: pCIG | Megason and McMahon, 2002 |  |
| Plasmid: pCAG-memRFP | This paper; Addgene | Addgene # 224564 |
| Plasmid: pCAG-Cas13d-T2A-Citrine | This paper; Addgene | Addgene # 224565 |
| Plasmid: pCAG-Cas13d-T2A-GFP(1-10) | This paper; Addgene | Addgene # 224566 |
| Plasmid: pCAG-memRFP-3xControlgRNA | This paper; Addgene | Addgene # 224567 |
| Plasmid: pCAG-nucGFP11-3xControlgRNA | This paper; Addgene | Addgene # 224568 |
| Plasmid: pCAG-3xGFP11-mem-3xControlgRNA | This paper; Addgene | Addgene # 224583 |
| Plasmid: pCAG-memRFP-3xPax7gRNA | This paper; Addgene | Addgene # 224569 |
| Plasmid: pCAG-nucGFP11-3xPax7gRNA | This paper; Addgene | Addgene # 224570 |
| Plasmid: pCAG-3xGFP11-mem-3xPax7gRNA | This paper; Addgene | Addgene # 224571 |
| Software and Algorithms | | |
| Fiji | Schindelin et al., 2012 | RRID:SCR_002285 |
| Zen 2 Blue | Zeiss |  |
| Photoshop CC | Adobe |  |
| Prism 10 | GraphPad |  |
